# Theta activity during encoding interacts with NREM sleep oscillations to predict memory generalization

**DOI:** 10.1101/2021.11.18.469190

**Authors:** Tamara Gibson, Zachariah R. Cross, Alex Chatburn

## Abstract

Relatively little is known regarding the interaction between encoding-related neural activity and sleep-based memory consolidation. One suggestion is that a function of encoding-related theta power may be to ‘tag’ memories for subsequent processing during sleep. This study aimed to extend previous work on the relationships between sleep spindles, slow oscillation-spindle coupling and task-related theta activity with a combined Deese-Roediger-McDermott (DRM) and nap paradigm. This allowed us to examine the influence of task- and sleep-related oscillatory activity on the recognition of both encoded list words and associative theme words. Thirty-three participants (29 females, mean age = 23.2 years) learned and recognised DRM lists separated by either a 2hr wake or sleep period. Mixed-effects modelling revealed the sleep condition endorsed more associative theme words and fewer list words in comparison to the wake group. Encoding-related theta power was also found to influence sleep spindle density, and this interaction was predictive of memory outcomes. The influence of encoding-related theta was specific to sleep spindle density, and did not appear to influence the strength of slow oscillation-spindle coupling as it relates to memory outcomes. The finding of interactions between wakeful and sleep oscillatory-related activity in promoting memory and learning has important implications for theoretical models of sleep-based memory consolidation.

---

One proposed function of episodic memory is to store information to allow accurate predictions about the environment. This is seen in particular through the established role of memory in influencing future behaviour, as has been shown in the relationship between episodic memory, long-term planning and imagination (Hassabis et al., 2007; Szpunar et al., 2013). It should be noted, however, that consolidation (and the accompanying generation of schemata) must necessarily rely on both encoded material and encoding processes, although how encoding-related brain activity may influence consolidation processes has rarely been investigated, at least in terms of related EEG factors.

Emerging work suggests that encoding-related theta activity is associated with successful memory consolidation (Hasselmo & Stern, 2014; Sederberg, Kahana, Howard, Donner & Madsen, 2003), and that increased theta activity during learning predicts higher post-learning sleep spindle density (Heib et al., 2015). There is a body of evidence for a role of encoding theta on memory outcomes: successful recall of encoded information has been shown to improve as a function of pre-stimulus theta amplitude (Cohen et al., 2015; Guderian et al., 2009; Heib et al., 2015). Similarly, artificially inducing theta oscillations with transcranial slow oscillation stimulation during the encoding of information results in better recall of encoded information (Kirov et al., 2009), and theta power increases during encoding predict successful consolidation of episodic memory (Sederberg et al., 2007). Together, this work suggests that encoding-related neural activity influences memory consolidation, although the specific relationship between theta power and neurophysiological aspects of sleep-based memory consolidation has not been thoroughly investigated. One perspective is that theta power at encoding may serve a “tagging” function for subsequent memory processing during sleep (Heib et al., 2015).

To date, research has largely ignored the role of encoding-related electrophysiological activity as an influencer of sleep-based memory consolidation. This study aimed to investigate the influences of encoding-related theta activity on elements of sleep-based memory consolidation, such as sleep spindles and slow oscillation-spindle coupling on both learning and associations using the DRM paradigm. Theta power was estimated during the learning phase of the DRM, which was followed by a two-hour retention period of either typical wakefulness or a nap. It was predicted that increased theta power at encoding would result in improved recognition of DRM list words and worsened performance for associations (i.e., endorsement of critical lures). It was also hypothesised that greater sleep spindle density would result in an increase in veridical memories. We also sought to replicate the relationship between encoding-related theta activity and sleep spindles reported by Heib at al. (2015), and to extend this relationship to determine whether encoding-related theta interacts with slow oscillation-spindle coupling to influence memory outcomes.

## Method

### Participants

Forty-three participants enquired, with 33 meeting the eligibility criteria. The final sample consisted of 33 participants between the ages of 18 and 33 (29 females, mean age = 23.2 years). Participants were randomly allocated into two groups, with 16 in the wake condition and 17 in the experimental nap condition. Sample size was determined through a G*Power calculation (Faul et al., 2007), which indicated that in order to obtain sufficient power (0.80) to detect a large effect size (0.80) at a significance level of .05, a sample of 30 participants was recommended (Diekelmann et al., 2008). All participants were healthy right-handed adults, who were not taking medication that could interfere with EEG, had not engaged in recreational drug use in the six months prior and were not diagnosed with a psychiatric or sleep disorder. The UniSA Human Research Ethics Committee granted ethics approval for the study, and all participants provided informed consent prior to participating. Participants received a $40 honorarium upon completion of the study.

### Screening measures

The PSQI was used to measure self-reported sleep quality in the month leading up to participation to screen for poor sleep quality (Buysse et al., 1989; Gobin et al., 2015). Participants with a score of 5 or above (maximum possible score of 21) were excluded from participating. The Finders Handedness Survey (Flanders) was also completed by participants as a self-report measure of hand preference. Left handers were excluded from participating to mitigate handedness-related differences in the EEG (Nicholls et al., 2013).

### Electroencephalography (EEG)

EEG data were collected using a BrainAmp BrainCap MR (Brain Products GmbH, Gilching, Germany), with 32-channel active DC Ag/AgCl electrodes. Electrodes were arranged according to the international 10-20 system (American Electroencephalographic Society, 1994). Bipolar electrooculogram (EOG) was also recorded, with electrodes placed 1cm diagonally from the outer canthus of each eye. The EEG was sampled at 1000 Hz with impedances for electrodes kept below 10kΩ. The online reference was located at FCz, and ground was AFz. EEG was continuously recorded during the DRM tasks, and during the sleep period.

### The Deese-Roediger-McDermott Paradigm (DRM)

The DRM was used as a semantic memory association task (Roediger & McDermott, 1995), and was presented using *OpenSesame* (Mathôt et al., 2012). Eighteen themed word lists were used in order to test for memory for specific items as well as production of the overall gist of the word-lists. The DRM lists have been reported to elicit associative memories 77% (mean of every list) of the time (Stadler et al., 1999). Split-half measurements of the DRM display high internal consistency (*r* = .80; Stadler et al., 1999).

Each list consisted of 14 study words and 2 critical lure words (not presented during the learning phase), with the order of the words during the learning task arranged from most to least related to the lures (Roediger & McDermott, 1995). To improve the quality and accuracy of EEG analysis, the first words from each list were added as a critical lure, in keeping with Beato & Diez (2011).

The learning phase involved presenting all 14 words from the 18 experimental study lists in a serial visual presentation format. Each word was presented for 1250ms. Each list word was followed by a blank screen for 250ms, then a fixation cross for 500ms. The learning phase took approximately 20 minutes. The recognition phases included the first, fifth and eighth word from each presented list and the same words from unrelated control lists. Two critical lure words were also included for recognition from each presented list and the control lists (See *Figure* 1a and 1b for a schematic of the learning and recognition phases, respectively). Unlike the learning phase, the words were presented pseudo-randomly, to ensure no words from the same list were presented in succession. The presentation format and timing in the recognition phase were identical to that used in the learning phase. Participants responded to the presented words based on whether they thought the words were presented during the learning phase (old) or not (new). This was performed with a keyboard press, and these responses allowed for four outcomes to be calculated for veridical and association memories: (1) hit; (2) false alarm; (3) miss, and; (4) correct rejection (see Jano et al., 2021 for more details).

**Figure 1.**
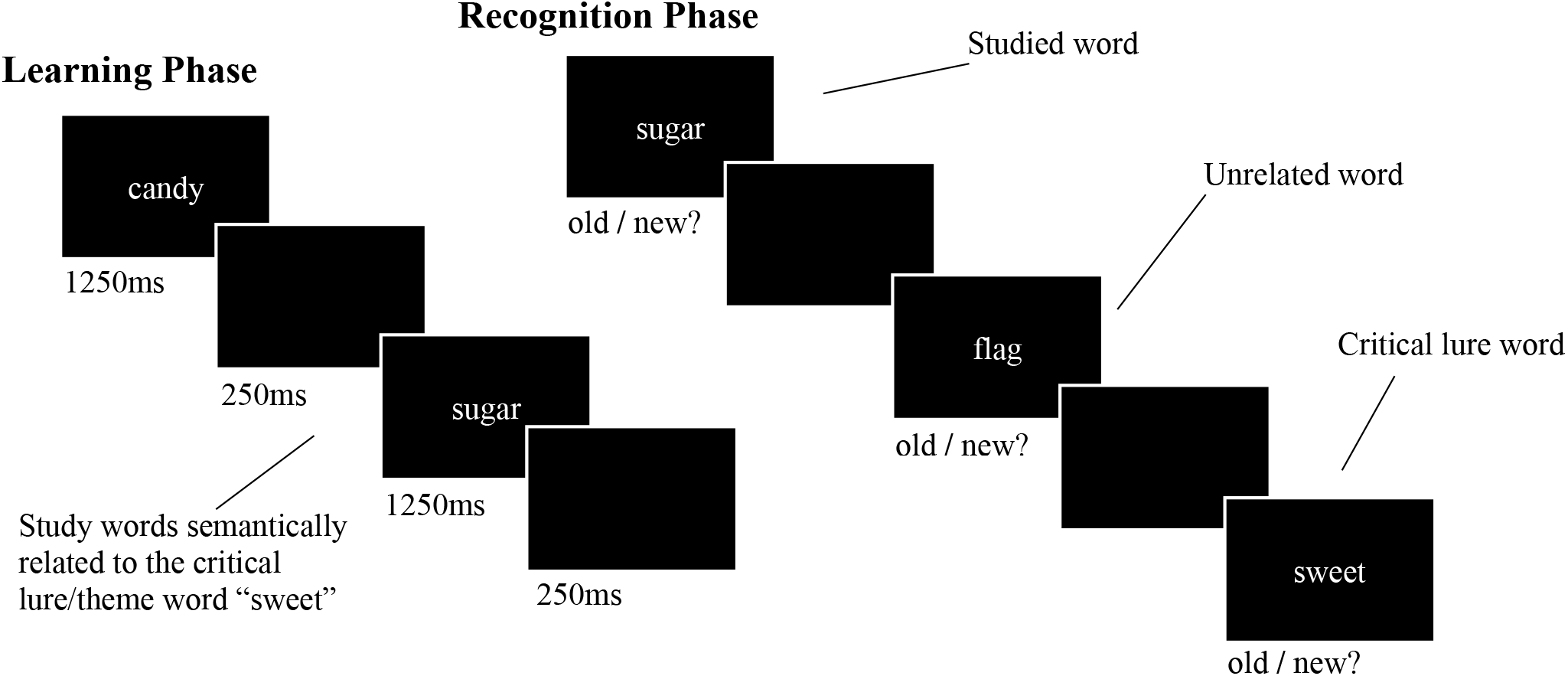
Example learning and recognition trials from the DRM paradigm. **a)** The learning phase involved viewing lists of related study words. **b)** The recognition phases involved immediate and delayed recognition, which involved the presentation of study words, their respective lures, and control words, to which participants made a self-paced old/new judgement.

### Procedure

Participants arrived at approximately 10:30am at the UniSA Cognitive and Systems Neuroscience-Research Hub and completed screening tools (the Flanders, the PSQI, and a demographic questionnaire). Participants were asked to reduce their sleep the night prior to testing by one hour to ensure participants experienced adequate sleep pressure for the afternoon nap, compliance of which was confirmed through self-report.

The EEG cap was then fitted, and participants were seated in a quiet testing room where the DRM paradigm was completed. EEG was recorded throughout the DRM task, and participants were instructed to do their best to commit list words to memory. Participants also undertook immediate recognition testing, in which a sub-list (three items) of each word list was presented to the participant, including the associative lures from each list. All conditions of stimuli presentation were consistent with the encoding phase. The same is true of the delayed recognition phase, in which the same content as in the immediate recognition and learning phases was presented in pseudo-randomised order.

Participants were randomly allocated to either the experimental (sleep), or to the control (wake) group. For the next two hours, participants either napped or stayed awake and were given free time to do quiet study or browse the internet (control; see *Figure 2*). The two-hour period was chosen to allow enough time for one sleep cycle. To align with the post-lunch circadian dip (Monk, 2005), the nap occurred at approximately 1:00pm.

**Figure 2.**
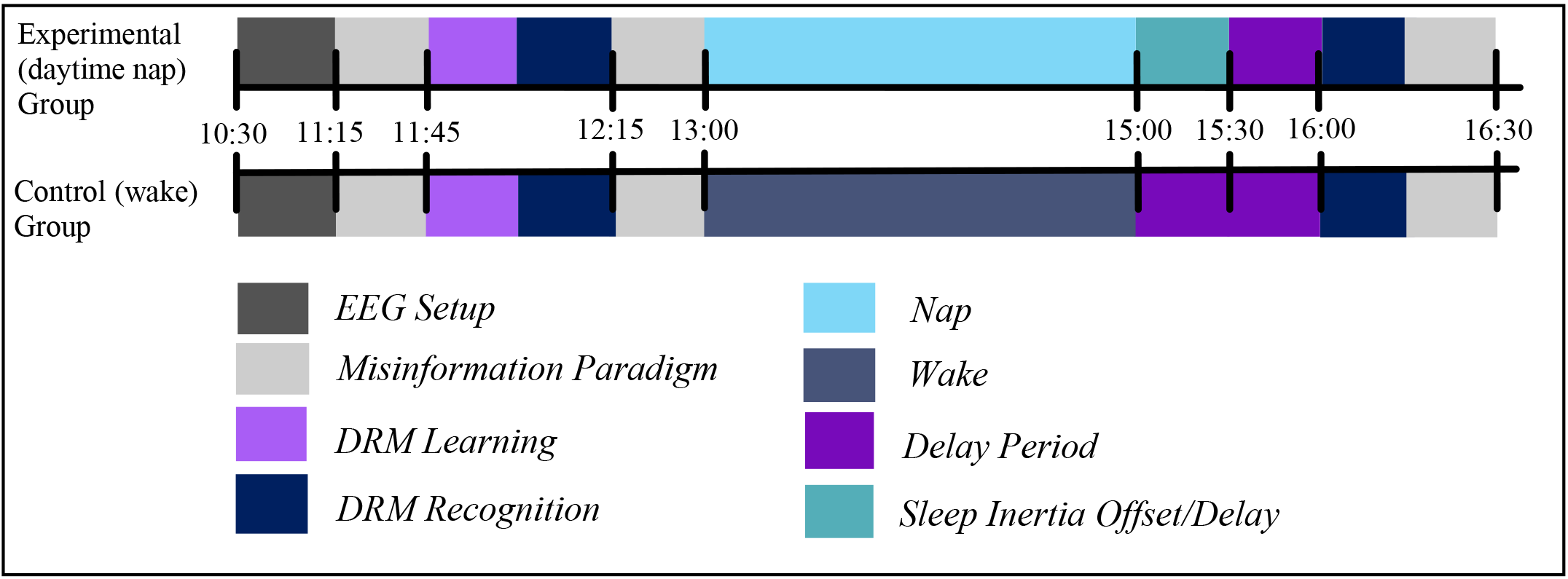
The experimental procedure. *Note.* Participants were fitted with EEG caps and undertook learning of DRM lists, before being tested for immediate recognition and, following a sleep/wake retention interval, delayed recognition. Note that both IR and DR periods involved the presentation of associative theme words. Also note the presence of a misinformation task, which was unrelated to the present analysis.

To compensate for sleep inertia (Achermann & Borbély, 1994), all participants were given a break of one hour following the nap. Participants then completed the delayed recognition (DR) phase, which involved presenting the recognition task of the DRM, which was the same task and stimuli in the IR, and participants’ EEG were again recorded. The recognition phase took approximately 15 minutes to complete.

## Data Analysis

### DRM Recognition Accuracy

Participant old/new responses were used to calculate d’ (McNicol, 2005) for both veridical and associative memories for both IR and DR tasks, as adapted from signal detection theory (McNicol, 2005). d’ was calculated using the package *Psycho* (Makowski, 2018) implemented in *R* version 3.6.1 (R Core Team, 2018). As per the methods used in Jano et al. (2021), d’ was calculated (Z_hits_ – Z_fa_) for both list words and associative lures, in order to reflect our viewpoint that ‘false memory’ measured in the DRM reflects associative processes, as opposed to memory errors.

### Sleep Scoring

An experienced technician followed the guidelines outlined by the American Academy of Sleep Medicine (AASM; Berry et al., 2012) to score participants’ sleep recordings in 30 second epochs. Information regarding total sleep time, sleep onset latency, and the percentage of time spent in sleep stages N1, N2, S3 and rapid eye movement (REM) sleep was acquired from this analysis.

### Sleep Spindle and Coupling Strength Detection

Sleep spindles were detected using the YASA toolbox implemented in MNE-Python (Vallat & Jajcay, 2020). The EEG signal was filtered between 12-16 Hz with a wide transition bandwidth of 1.5 Hz. The amplitude was calculated by applying a Hilbert transform which was then smoothed with a 200ms moving average. Candidate EEG phenomena which exceeded calculated thresholds were considered spindles, where the length of the spindle was determined by the length of time that absolute sigma power and covariance exceed their threshold. If the identified event was <0.3s or >2.5s, it was rejected. Slow oscillation-spindle coupling strength was also detected using the same toolbox, which was based on published algorithms (Helfrich et al., 2018). Slow oscillations were first extracted through continuous NREM EEG data and filtered using a digital phase-true FIR band-pass filter from 0.3 – 2 Hz with a 0.2 Hz transition band to detect zero crossing events that were between 0.3 – 1.5 s in length, and that met a 75 to 500 microvolt criterion. Slow oscillation-spindle coupling was detected using an event-locked cross frequency coupling metric. The normalised slow oscillation trough-locked data was filtered into the slow oscillation component (0.1 – 1.25 Hz) and extracted the instantaneous phase angle after applying a Hilbert transform. The same trials were then filtered between 12 – 16 Hz and then the instantaneous amplitude from the Hilbert transform was extracted. For each participant at channel Cz and epoch, the maximal sleep spindle amplitude and corresponding slow oscillation phase angle was calculated. The mean resultant vector length (mean vector length; coupling strength) across all NREM events was then determined using circular statistics implemented in the *pingouin* package (Vallat, 2018). A scale of 0 – 1 was used for mean vector length, with 1 indicating that each coupled spindle occurred at the same phase of the slow oscillation, and 0 indicating that each coupled spindle occurred at a different phase of the slow oscillation.

### DRM Time Frequency Analysis

Spectral activity in the theta range (~ 4 – 7 Hz) was estimated using a complex Morlet wavelet analysis implemented in MNE-Python (tfr_morlet). Participants’ theta frequency ranges were adjusted according to the golden mean algorithm (Klimesch, 2012). Theta power was estimated from -200 to 1000ms post-word onset during the immediate and delayed DRM tasks.

### Linear Mixed-Effects Models

Linear mixed effects models were conducted using the *lme4* package (Bates at al., 2014) in *R* (R Core Team, 2018) to assess the interactions between encoding-related theta power and consolidation-related sleep physiology on the generation of veridical and association memories.

To test hypothesis one (theta power increases at encoding would lead to an increase in memory for DRM list words and a decrease in endorsement of associative theme words), we aimed to predict d’ from task-evoked theta power, condition (sleep/wake) and memory type (veridical/association), while controlling for the effects of immediate recognition testing performance. Models also included mean pre-stimulus theta power as a scaled covariate in order to control for pre-stimulus activity (Alday, 2018), and by-channel random intercepts to account for topographical differences in theta power estimates. By-participant random intercepts were also specified in each model.

To explore our second hypothesis (greater sleep spindle density would result in an increase in memory for DRM list words), as well as our research questions around the role of encoding-related theta power as a driver of both sleep spindle density as well as association of list words across the retention period, encoding-related theta power, sleep spindle density and memory type were used as predictors of d’ from the pre- to post-retention intervals in the sleep condition, once again, controlling for immediate recognition performance.

To test whether encoding-related theta power interacts with slow oscillation-spindle coupling strength to predict memory outcomes, we modelled d’ scores across retention period from theta power at encoding, memory type, slow oscillation-spindle coupling strength, from participants in the sleep condition.

## Results

d’ scores per condition are reported in Table 2. The average associative memory endorsement rate across participants, conditions, and recognition period was 67.66%, demonstrating successful DRM effects. Sleep parameters from the experimental condition are reported in Table 3

**Table 2.** Mean (SD) d’ Scores Across Task, Condition and Memory Type.

| Condition | Immediate recognition |  | Delayed recognition |  |
| --- | --- | --- | --- | --- |
|  | List words | Theme words | List words | Theme words |
| Sleep | 1.58 (0.64) | 1.24 (0.37) | 1.28 (0.64) | 1.28 (0.50) |
| Wake | 1.50 (0.43) | 1.22 (0.43) | 1.14 (0.44) | 1.13 (0.55) |

**Table 3.**
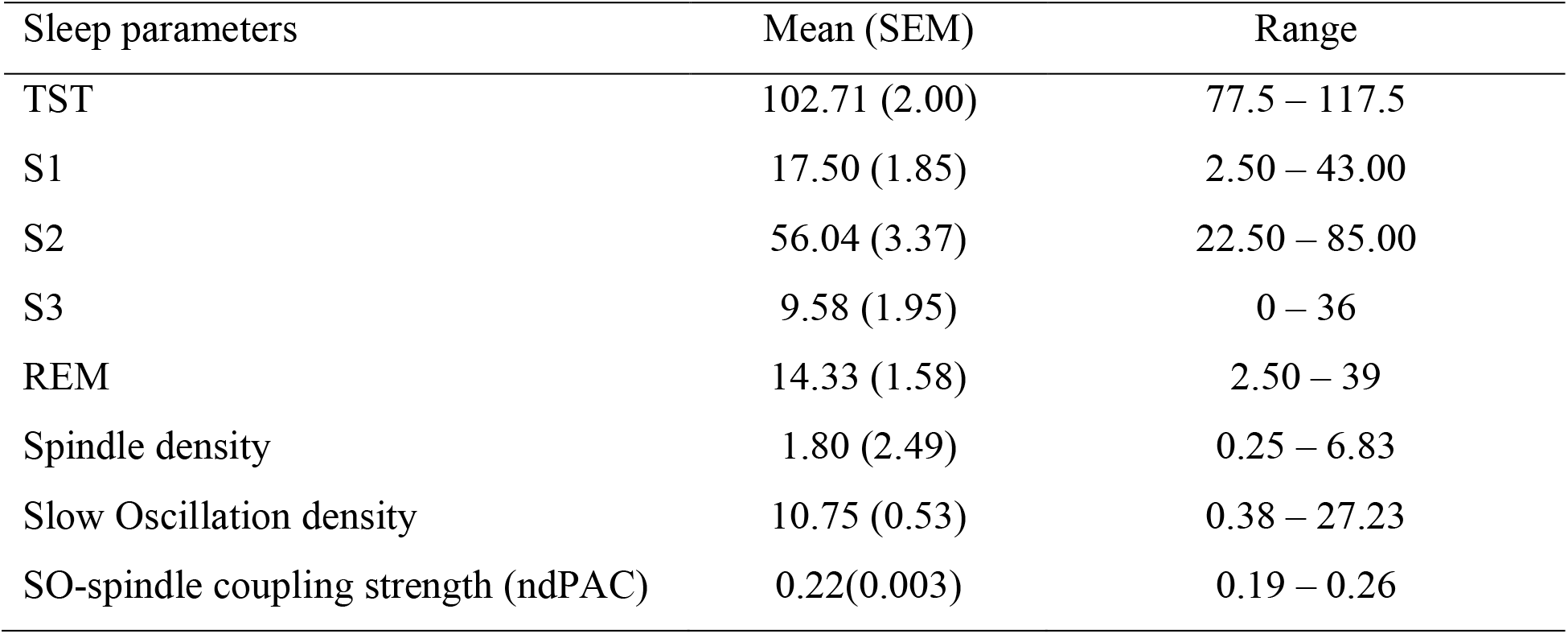
Time Spent Asleep, Sleep Spindle, Slow Oscillation and Spindle\SO Coupling Strength as Means, Standard Errors and Ranges.

Behavioural results indicate a significant relationship between condition (sleep/wake) and type of memory (list word or associative lure) in determining memory for list words and associative lures (χ2(1) = 13.65, *p* <.001), such that associative lures were recognised with slightly greater accuracy than list words (see figure 3A). Testing of hypothesis 1 indicated a significant theta × condition × memory interaction (χ2(1) = 43.86, *p* <.001), such that relative theta power increases at encoding resulted in the recognition of fewer list word memories for the wake condition but greater successful recognition for the sleep condition. This analysis demonstrated the opposite relationship for the endorsement of theme words (i.e., associations): relative theta decreases at encoding led to more endorsement of theme words and lower recognition of list words in the context of sleep (see Figure 3B).

**Figure 3.**
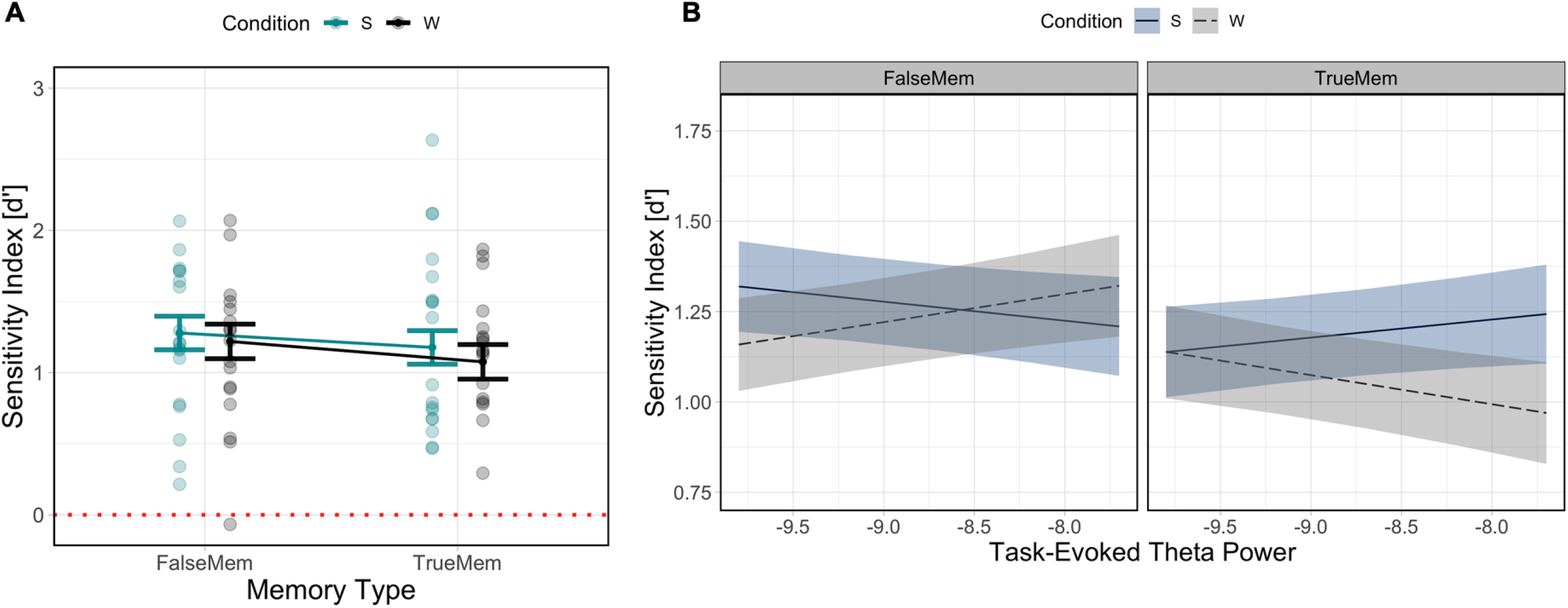
Behavioural (A) and power spectral (B) results in determining memory for list words and associative lures. In A) behavioural results indicate a slight increase in accuracy in the recognition of associative lures across sleep and wake retention intervals. In B) results indicate a complex relationship between theta power at encoding, memory type and sleep/wake. Dashed line indicates chance performance.

Statistical analysis to test hypothesis 2 indicated a significant interaction was found between theta power and spindle density in determining memory for list words and associative lures, such that theta increases in conjunction with lower spindle density was associated with an increase in theme word recognition and a decrease in list word recognition (χ2(1) = 15.51, *p* <.001). This relationship was modulated as a function of sleep spindle density (see Figure 4A). Similarly, we observed a significant interaction between slow oscillation-spindle coupling, memory type and theta power in determining outcomes (χ2(1) = 7.12, *p* = .008), such that theta power increases were differentially associated with memory for list words and lures across coupling strength levels (see Figure 4B). List word recognition was not modulated as a function of the interaction between slow oscillation-spindle coupling and theta power.

**Figure 4.**
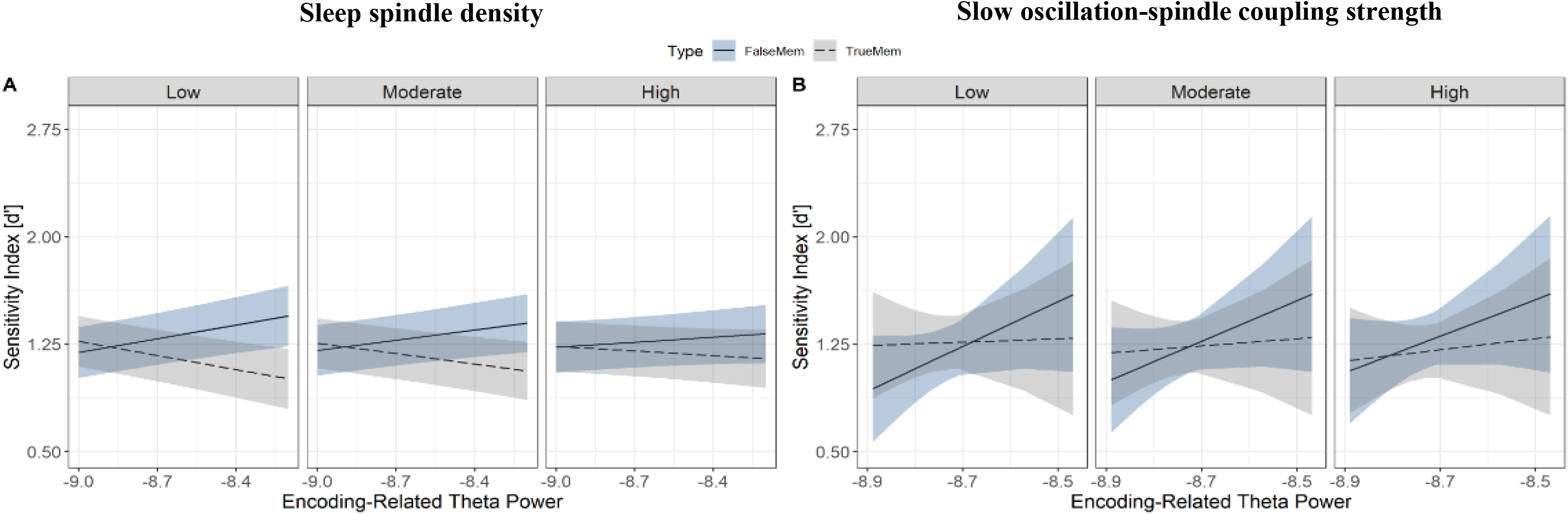
Relationships between sleep microstate variables and memory for list words and associative lures. *Notes.* (A) Interaction between theta power at encoding (x-axis), memory type (association in blue, veridical in grey), and sleep spindle density (facets) in predicting d’(y-axis); (B) Interaction between theta power at encoding (x-axis), memory type (association in blue, veridical in grey), and sleep spindle-slow oscillation coupling strength (facets) in predicting d’ performance (y-axis). Ribbons indicate the 83% confidence interval.

## Discussion

Here, we examined how encoding-related theta activity influences memory consolidation across sleep to facilitate learning and memory. We sought to test the ideas that theta power at encoding would influence recognition of DRM list and theme words, potentially through the modulation of sleep-related oscillatory microstructure. To this end, we tested the relationship between encoding-related theta power and both sleep spindle density and slow oscillation-spindle coupling strength. Contrary to the first hypothesis (that relative theta power increases would lead to an increase in recognition of list words and a decrease in endorsement of associative theme words) we note an effect of relative theta decreases which differentially impacted memory between the wake and sleep conditions: wake resulted in improved recognition of list words and a decrease in endorsement of associative lures, both of which were tracked by encoding-related theta. Sleep was related to a decrease in recognition of list words and an increase in theme word endorsement, both linked to relative theta power decreases. This suggests that theta at encoding relates to both memory for experienced episodes and the association thereof and that this is differentially modulated by sleep and wake. As such, our results expand previous work on the role of encoding-related brain activity and how it may relate to sleep-based consolidation mechanisms.

Theta power during the encoding of information appears to play an important role in subsequent memory outcomes (Hasselmo & Stern, 2014; Sederberg et al., 2003). Our results indicate that theta power decreases at encoding differentially effects both recognition of previously seen, as well as gist memories, and has differential effects as a function of wake versus sleep. These results are conceptually similar, but directionally different to those published in previous EEG studies, which have found that theta power increases, not decreases, relates to successful encoding of episodic memories (Heib et al., 2015; Kirov et al., 2009). This difference in findings can be explained by methodological differences between our study and previous literature. We have used a more direct measurement of theta power during encoding, whereas Heib and colleagues (2015) used a calculated difference between immediate recall testing and a non-memory section of their paradigm in order to determine encoding-related theta activity. The cognitive implications of this difference may be that we have measured brain activity related to encoding a memory trace, whereas Heib et al. may have measured activity related to reconsolidation. Whether or not this is a meaningful difference is presently unclear and should be the topic of future enquiry.

Previous literature (Heib et al., 2015; but more so Kirov et al., 2009) elegantly demonstrates that encoding theta power relates to memory outcomes across sleep. Typically, these findings are explained through an influence on hippocampocortical communication at theta frequency during wakeful encoding. Our study is unable to comment on whether this is specifically a ‘tagging’ mechanism, as we did not include a manipulation to differentiate between memories to be consolidated and those to be ignored. Despite this, our results can still be interpreted through similar mechanisms as previously published, through the idea that theta may prime circuits for subsequent consolidation during sleep, and thus increase sleep spindle activity at these local sites during post-learning sleep (Petzka, Chatburn, Charest, Balanos & Staresina, 2021).

Our findings also demonstrate a clear relationship between encoding-related theta power and sleep spindle density. This partially supports the findings of Heib et al. (2015), with a difference of the directionality of theta power. These results further support the proposed effect of theta oscillations on consolidation through modulation of sleep spindle density (Heib et al., 2015). Encoding-related theta may potentially be an important marker of relevant encoding-related activity, which serves to prime circuits for subsequent consolidation, in much the same way that topographical overlap between encoding-related brain activity and sleep spindle density may relate to consolidation and memory outcomes across sleep (Petzka et al., 2021). Previous research can be updated based on this, mainly looking at a sleep spindle specific role in the consolidation of different types of memory (Smith, et al., 2007), such that gist processing is supported via higher spindle density along with theta power increases at encoding, and recognition of previously seen items showing the opposite trend. Sleep spindles have been implicated with integrating new information into existing knowledge (Tamminen et al., 2010), suggesting that their function reflects a more general process of learning as well as the hippocampocortical communication of encoded memory traces. It may be informative to consider the mechanisms through which encoding-related theta power and spindle density in subsequent sleep may lead to both improvements in memory, and the generalisation of encoded traces. Both mechanisms could be accounted for through statistical regularities in encoded information leading to greater replay for shared components over individual components, as is theorised in the competitive trace and IoTa accounts of memory (Yassa & Reagh, 2013; Lewis & Durrant, 2011). This idea has also been examined in memory literature to explain attributes of memory such as selective consolidation, item integration and associations, along with how consolidation binds that information together (Stickgold & Walker, 2013; Walker & Stickgold, 2010). A common mechanism to explain the capacity of the human brain to both use pattern separation processes to store veridical aspects of our experience, as well as to use pattern completion processes to extract meanings and rules therefrom would seem to be efficient. Our finding that different polarities of theta power at encoding may predict outcomes in terms of both memory and learning should be investigated more fully in future research, potentially through causal manipulations using associative memory paradigms, naps and non-invasive brain stimulation techniques, or different training modalities.

There are several limitations to the present study which should be considered in the interpretation of results. Considering that the DRM is fairly artificial, the generalisability of these findings should be tempered Additionally, the DRM scores observed are higher than noted in other DRM studies (Pardilla-Delgado & Payne, 2017), which may be due to the shorter time frame between encoding and recognition, along with the effect of including more trials and an additional critical lure. Additional, nonspecific effects such as this may have influenced participants in their encoding and generalisation of information. This should not exert a marked influence on the EEG, although a more standard and controlled behavioural procedure would be of benefit in fortifying the present results. Further, considering that sleep loss prior to learning can negatively impact memory ability (Alberca-Reina et al., 2015; Mander et al., 2008), our manipulation of requiring participants to restrict their sleep by one hour prior to encoding may have influenced their capacity to encode the material, although it is unlikely that any effects of this procedure were of a significant nature.

Research has also suggested that the aperiodic activity directly following a spindle may also be important for consolidation (Helfrich et al., 2021). The authors suggest the role of a sleep spindle is similar to that of being a messenger signal rather than the producer of consolidation itself, as is currently theorised. That is, by not analysing aperiodic activity around sleep spindles, we may have missed an important factor in sleep-based memory consolidation. This is a clear area of importance for future research.

With the idea that theta acts a tagging mechanism, the relationships found in the present study demonstrates that encoding-related theta activity can influence specific aspects of memory-related sleep neurophysiology, and highlights the connection between encoding and consolidation process in episodic memory. Consolidation is clearly impacted by processes tagged by theta power during encoding, and if models wish to develop a comprehensive view of memory consolidation, then extant theories of sleep-based memory consolidation should expand to include the neurobiological mechanisms underlying successful memory encoding.

## Acknowledgements

The authors would like to thank Matthias Schlesewsky for insightful discussions during study design

## References

Achermann, P., & Borbély, A. A. (1994). Simulation of daytime vigilance by the additive interaction of a homeostatic and a circadian process. Biological Cybernetics, 71(2), 115–121.

Alberca-Reina, E., Cantero, J. L., & Atienza, M. (2015). Impact of sleep loss before learning on cortical dynamics during memory retrieval. NeuroImage, 123, 51–62. https://doi.org/10.1016/j.neuroimage.2015.08.033

Beato, M. S., & Diez, E. (2011). False recognition production indexes in Spanish for 60 DRM lists with three critical words. Behavior Research Methods, 43(2), 499–507. https://doi.org/10.3758/s13428-010-0045-9

Berry, R. B., Brooks, R., Gamaldo, C. E., Harding, S. M., Marcus, C., & Vaughn, B. V. (2012). The AASM manual for the scoring of sleep and associated events. Rules, Terminology and Technical Specifications, Darien, Illinois, American Academy of Sleep Medicine, 176, 2012.

Buysse, D. J., Reynolds, C. F., Monk, T. H., Berman, S. R., & Kupfer, D. J. (1989). The Pittsburgh Sleep Quality Index: A new instrument for psychiatric practice and research. Psychiatry Res, 28(2), 193–213.

Cohen, N., Pell, L., Edelson, M. G., Ben-Yakov, A., Pine, A., & Dudai, Y. (2015). Peri-encoding predictors of memory encoding and consolidation. Neuroscience & Biobehavioral Reviews, 50, 128–142. https://doi.org/10.1016/j.neubiorev.2014.11.002

Cross, Z. R., Helfrich, R. F., Corcoran, A. W., Kohler, M. J., Coussens, S., Zou-Williams, L., Schlesewsky, M., Gaskell, M. G., Knight, R. T., & Bornkessel-Schlesewsky, I. (2021). Spindle-slow oscillation coupling during sleep predicts sequence-based language learning. BioRxiv, 2020.02.13.948539. https://doi.org/10.1101/2020.02.13.948539

Diekelmann, S., Landolt, H. P., Lahl, O., Born, J., & Wagner, U. (2008). Sleep loss produces false memories. PLoS ONE [Electronic Resource], 3(10), e3512. MEDLINE. https://doi.org/10.1371/journal.pone.0003512

Faul, F., Erdfelder, E., Lang, A.-G., & Buchner, A. (2007). G*Power 3: A flexible statistical power analysis program for the social, behavioral, and biomedical sciences. Behavior Research Methods, 39(2), 175–191. https://doi.org/10.3758/BF03193146

Gobin, C. M., Banks, J. B., Fins, A. I., & Tartar, J. L. (2015). Poor sleep quality is associated with a negative cognitive bias and decreased sustained attention. Journal of Sleep Research, 24(5), 535–542.

Gramfort, A., Luessi, M., Larson, E., Engemann, D. A., Strohmeier, D., Brodbeck, C., Goj, R., Jas, M., Brooks, T., Parkkonen, L., & Hämäläinen, M. (2013). MEG and EEG data analysis with MNE-Python. Frontiers in Neuroscience, 7. https://doi.org/10.3389/fnins.2013.00267

Guderian, S., Schott, B. H., Richardson-Klavehn, A., & Duzel, E. (2009). Medial temporal theta state before an event predicts episodic encoding success in humans. Proceedings of the National Academy of Sciences, 106(13), 5365–5370. https://doi.org/10.1073/pnas.0900289106

Hassabis, D., Kumaran, D., Vann, S. D., & Maguire, E. A. (2007). Patients with hippocampal amnesia cannot imagine new experiences. Proceedings of the National Academy of Sciences, 104(5), 1726. https://doi.org/10.1073/pnas.0610561104

Hasselmo, M. E., & Stern, C. E. (2014). Theta rhythm and the encoding and retrieval of space and time. Neuroimage, 85, 656–666.

Heib, D. P. J., Hoedlmoser, K., Anderer, P., Gruber, G., Zeitlhofer, J., & Schabus, M. (2015). Oscillatory Theta Activity during Memory Formation and Its Impact on Overnight Consolidation: A Missing Link? Journal of Cognitive Neuroscience, 27(8), 1648–1658. https://doi.org/10.1162/jocn_a_00804

Helfrich, R. F., Lendner, J. D., & Knight, R. T. (2021). Aperiodic sleep networks promote memory consolidation. Trends in Cognitive Sciences, 25(8), 648–659. https://doi.org/10.1016/j.tics.2021.04.009

Helfrich, R. F., Mander, B. A., Jagust, W. J., Knight, R. T., & Walker, M. P. (2018). Old Brains Come Uncoupled in Sleep: Slow Wave-Spindle Synchrony, Brain Atrophy, and Forgetting. Neuron, 97(1), 221–230.e4. https://doi.org/10.1016/j.neuron.2017.11.020

Jano, S., Romeo, J., Hendrickx, M. D., Schlesewsky, M., & Chatburn, A. (2021). Sleep influences neural representations of true and false memories: An event-related potential study. Neurobiology of Learning and Memory, 107553.

Kirov, R., Weiss, C., Siebner, H. R., Born, J., & Marshall, L. (2009). Slow oscillation electrical brain stimulation during waking promotes EEG theta activity and memory encoding. Proceedings of the National Academy of Sciences, 106(36), 15460–15465. https://doi.org/10.1073/pnas.0904438106

Klimesch, W. (2012). Alpha-band oscillations, attention, and controlled access to stored information. Trends in cognitive sciences, 16(12), 606–617.

Lewis, P. A., & Durrant, S. J. (2011). Overlapping memory replay during sleep builds cognitive schemata. Trends in cognitive sciences, 15(8), 343–351.

Makowski, D. (2018). The psycho package: An efficient and publishing-oriented workflow for psychological science. Journal of Open Source Software, 3(22), 470.

Mander, B. A., Reid, K. J., Davuluri, V. K., Small, D. M., Parrish, T. B., Mesulam, M.-M., Zee, P. C., & Gitelman, D. R. (2008). Sleep deprivation alters functioning within the neural network underlying the covert orienting of attention. Brain Research, 1217, 148–156. https://doi.org/10.1016/j.brainres.2008.04.030

Mathôt, S., Schreij, D., & Theeuwes, J. (2012). OpenSesame: An open-source, graphical experiment builder for the social sciences. Behavior Research Methods, 44(2), 314–324.

McNicol, D. (2005). A primer of signal detection theory. Psychology Press.

Monk, T. H. (2005). The post-lunch dip in performance. Clinics in Sports Medicine, 24(2), e15–e23.

Nicholls, M. E., Thomas, N. A., Loetscher, T., & Grimshaw, G. M. (2013). The Flinders Handedness survey (FLANDERS): A brief measure of skilled hand preference. Cortex, 49(10), 2914–2926.

Petzka, M., Chatburn, A., Charest, I., Balanos, G. M., & Staresina, B. P. (2021). Sleep spindles track cortical learning patterns for memory consolidation. bioRxiv.

Roediger, H. L., & McDermott, K. B. (1995). Creating false memories: Remembering words not presented in lists. Journal of Experimental Psychology: Learning, Memory, and Cognition, 21(4), 803.

Sederberg, P. B., Kahana, M. J., Howard, M. W., Donner, E. J., & Madsen, J. R. (2003). Theta and gamma oscillations during encoding predict subsequent recall. Journal of Neuroscience, 23(34), 10809–10814.

Sederberg, P. B., Schulze-Bonhage, A., Madsen, J. R., Bromfield, E. B., Litt, B., Brandt, A., & Kahana, M. J. (2007). Gamma Oscillations Distinguish True From False Memories. Psychological Science, 18(11), 927–932. https://doi.org/10.1111/j.1467-9280.2007.02003.x

Stadler, M. A., Roediger, H. L., & McDermott, K. B. (1999). Norms for word lists that create false memories. Memory & Cognition, 27(3), 494–500.

Stickgold, R., & Walker, M. P. (2013). Sleep-dependent memory triage: Evolving generalization through selective processing. Nature Neuroscience, 16(2), 139–145. https://doi.org/10.1038/nn.3303

Szpunar, K. K., Addis, D. R., McLelland, V. C., & Schacter, D. L. (2013). Memories of the Future: New Insights into the Adaptive Value of Episodic Memory. Frontiers in Behavioral Neuroscience, 7. https://doi.org/10.3389/fnbeh.2013.00047

Tamminen, J., Payne, J. D., Stickgold, R., Wamsley, E. J., & Gaskell, M. G. (2010). Sleep Spindle Activity is Associated with the Integration of New Memories and Existing Knowledge. Journal of Neuroscience, 30(43), 14356–14360. https://doi.org/10.1523/JNEUROSCI.3028-10.2010

Vallat, R., & Jajcay, N. (2020). raphaelvallat/yasa: V0.3.0. Zenodo. https://doi.org/10.5281/zenodo.3818479

Vallat, R. (2018). Pingouin: Statistics in Python. Journal of Open Source Software, 3(31), 1026. https://doi.org/10.21105/joss.01026

Walker, M. P., & Stickgold, R. (2010). Overnight alchemy: Sleep-dependent memory evolution. Nature Reviews Neuroscience, 11(3), 218–218. https://doi.org/10.1038/nrn2762-c1

Wilhelm, I., Diekelmann, S., Molzow, I., Ayoub, A., Molle, M., & Born, J. (2011). Sleep Selectively Enhances Memory Expected to Be of Future Relevance. Journal of Neuroscience, 31(5), 1563–1569. https://doi.org/10.1523/JNEUROSCI.3575-10.2011

Yassa, M. A., & Reagh, Z. M. (2013). Competitive trace theory: a role for the hippocampus in contextual interference during retrieval. Frontiers in behavioral neuroscience, 7, 107.

